# Function of the *Vibrio* PomB plug region based on the stator rotation model for bacterial flagellar motor

**DOI:** 10.1101/2021.03.16.435749

**Authors:** Michio Homma, Hiroyuki Terashima, Hiroaki Koiwa, Seiji Kojima

## Abstract

Bacterial flagella are the only real rotational motor organs in the biological world. The spiral-shaped flagellar filaments that extend from the cell surface rotate like a screw to create a propulsive force. The base of the flagellar filament has a protein motor consisting of a stator and a rotor embedded in the membrane. The motor part has stators composed of two types of membrane subunits, PomA(MotA) and PomB(MotB), which are energy converters coupled to the ion flow that assemble around the rotor. Recently, structures of the stator, in which two molecules of MotB stuck in the center of the MotA ring made of five molecules, were reported and a model in which the MotA ring rotates with respect to MotB, which is coupled to the influx of ions, was proposed. We focused on the *Vibrio* PomB plug region, which has been reported to control the activation of flagellar motors. We searched for the plug region, which is the interacting region, through site-directed photo-cross-linking and disulfide cross-linking experiments. Our results demonstrated that it interacts with the extracellular short loop region of PomA, which is between transmembrane 3 and 4. Although the motor halted following cross-linking, its function was recovered with a reducing reagent that disrupted the disulfide bond. Our results support the hypothesis, which has been inferred from the stator structure, that the plug region terminates the ion inflow by stopping the rotation of the rotor.

**Importance:** The flagellar biological motor resembles a mechanical motor, which is composed of stator and rotor and where the rotational force is transmitted by gear-like movements. We hypothesized that the flagellar the rotation of stator that the pentamer of A subunits revolves around the axis of the B subunit dimer with ion flow. The plug region of the B subunit has been shown to regulate the ion flow. Herein, we demonstrated that the ion flow was terminated by the crosslinking between the plug region and the A subunit. These finding support the rotation hypothesis and explain the role of the plug region in terminating the rotation.

## Introduction

F-type ATP synthase, V/A-type ATP enzyme, and bacterial flagella are well-known molecular motors driven by ion-driven forces (1). Bacterial flagella, besides spirochete, generate the driving force by rotating filaments with a structure similar to spiral microtubules extending from the cell surface as screws. The rotational force is generated by the interaction between the rotor, which is the cytoplasmic C-ring of the flagellar basal body, and the stator, which is a membrane protein complex that forms an ion transporter. The rotor is attached to a transmembrane MS-ring on top of the C-ring and is further connected to the extracellular filament via structures, such as rods and hooks. Therefore, when the rotor rotates, the filament portion rotates. In many bacteria, the C-ring is composed of three types of proteins: FliG, FliM, and FliN. Mutations in these proteins result in defects with several phenotypes, such as flagellar formation, motor rotation, or switching between CCW and CW rotations (2, 3). The genotypes of the phenotypes are indicated by *fla*, *mot*, and *che* (4). The C-terminal region of FliG is thought to interact with the cytoplasmic region of the stator to generate torque (1). MotA and MotB of *E. coli*, and PomA and PomB of *Vibrio* spp. are membrane proteins found in the stator complex. The A subunit has four transmembrane segments (TMs) and a large cytoplasmic region between TM2 and TM3. The B subunit has one TM in which aspartic acid (D), an essential ion bonding residue, is present in its N-terminal region and has a peptidoglycan binding (PGB) domain in its C-terminal region.

The polar flagellum of *Vibrio* spp. is driven by sodium-motive force (SMF), while the peritrichous flagellum of *E. coli* is driven by the proton-motive force (PMF). The motility of *Vibrio* is specifically inhibited by amiloride and its derivatives, which are inhibitors of epidermal sodium channels in animals (5, 6). The mutations conferring resistance to the amiloride derivative, phenamil, are mapped to the TM region of PomB and the TM3 region of PomA at the site of the cytoplasmic side membrane interface, and the vicinity is considered a part of the sodium ion transport pathway (7, 8). *Vibrio* motor proteins were initially thought to consist of the single transmembrane proteins, MotX and MotY, which are very different from the proton-type motor proteins (9–11). However, a new gene complementing the *mot* phenotype was identified and named the polar flagellar motors, *pomA* and *pomB* (12). Sequences of *pomA* and *pomB* are not highly homologous to that of the proton motor genes, *motA* and *motB*, but are structurally distinct as four transmembrane and one transmembrane proteins, and have homologous key residues. The transmembrane regions of MotX and MotY are the signal sequences for cell membrane transport. In fact, they are known to exist as rings in the outer membrane and are responsible for anchoring the stator complex (13, 14).

The ion-transporting stator complexes are thought to assemble at least 11 units around the rotor and interact with the FliG of the C-ring (15). They are anchored by the interaction of domains with extracellular PGB motifs of PomB or MotB (16, 17). Once incorporated, the stator complex allows ions to flow and motors to rotate. Moreover, it is believed that structural changes induced by the interaction between the cytoplasmic region of PomA and the C-terminal region of the rotator protein FliG activate the stator (18) and a large conformational change is speculated to occur in the periplasmic region at that time (19).

The inactivation of the stator is attributed to the plug region of the B subunit in *E. coli* (20). Deletion of the region from amino acid residues 51 to 70 immediately after the transmembrane region of *E. coli* MotB and its expression from the plasmid with MotA led to growth inhibition. Even in PomB, when the linker region containing the plug of 80 residues was deleted from E41 of PomB immediately after the transmembrane region, growth was inhibited with PomA; however, motility was not lost (21). Protons and sodium ions flow into the cells due to the expression of the stator lacking the plug region (22, 23). However, how the plug region blocks the influx of ions remains unknown.

The A and B subunits, which form the stator complex of the flagellar motor, are not highly homologous in sequence, but are functionally highly compatible. When PotB, a chimera between the C-terminal region of MotB and the N-terminal region of PomB, is expressed with PomA in *E. coli*, the *E. coli* flagella, which are originally driven by protons, are driven by sodium ions (24 49). Conversely, when *E. coli* MotAB is introduced into *Vibrio* cells, a very poor motility was exhibited. Furthermore, *Vibrio* motility can be conferred to the flagellum without MotXY, which is required for PomAB. Surprisingly, the MotB chimera between the N-terminal region of *A. aeolicus* MotB and the C-terminal periplasmic region of *E. coli* was expressed with *A. aeolicus* MotA in *E. coli*, which could function in *E. coli* as a sodium-driven motor (25). Such finding implies that the robustness of the rotor-stator interaction is very high.

Structural information is indispensable to understand the mechanism of force generation in bacterial flagellar motors. Until recently, only low-resolution density maps of stator units obtained by single-particle analysis using electron microscopy were available (25, 26). However, the atomic structures of MotA/MotB derived from *Campylobacter jejuni*, *Clostridium sporogenes*, and *Bacillus subtilis* have been reported in two different groups (27, 28). The MotA/MotB and PomA/PomB complexes, previously proposed as 4:2 heterohexamers, were shown to be 5:2 heteroheptamers in the present analysis. Based on the structure, in which two molecules of MotB are inserted in the center of the MotA ring comprised of five molecules, a model in which the MotA ring rotates with respect to the MotB axis due to the ion influx was proposed. In this study, we focused on the plug region of *Vibrio* PomB, which reportedly regulates the activation of the flagellar motor, and searched for regions of PomA that interact with the plug region using a site-specific *in vivo* photo-crosslinking technology. The purpose of this study was to clarify how the plug region regulates the ion flux.

## Results

### Detection of photo-crosslinked products of the PomB plug region and PomA

As the *Vibrio* cells used in this study are resistant to ampicillin, various mutations were introduced into pHFAB, which was cloned into a pBAD33-based chloramphenicol-resistant plasmid that can be expressed in both *E. coli* and *Vibrio* for functional analysis. As the vector for detecting protein-protein interactions by photo-crosslinking, which contains the amber suppressor tRNA, is on a chloramphenicol-resistant plasmid, we cloned the *pomAB* genes into the ampicillin-resistant pBAD24 plasmid. The *pomAB* genes in which an amber codon was introduced into the gene corresponding to residues F47 to A57 of PomB, were expressed in *E. coli* strain DH5α (Fig. 1). The amber suppressor tRNA was charged with pBPA, a phenylalanine derivative, for protein synthesis. By photo-crosslinking with UV, we detected bands with increased molecular weight in F47, I50, A51, and M54, which were detected by the anti-PomA as well as anti-PomB antibodies (Fig. 2). These bands indicated crosslinked products between the PomA and PomB molecules.

**Fig. 1.**
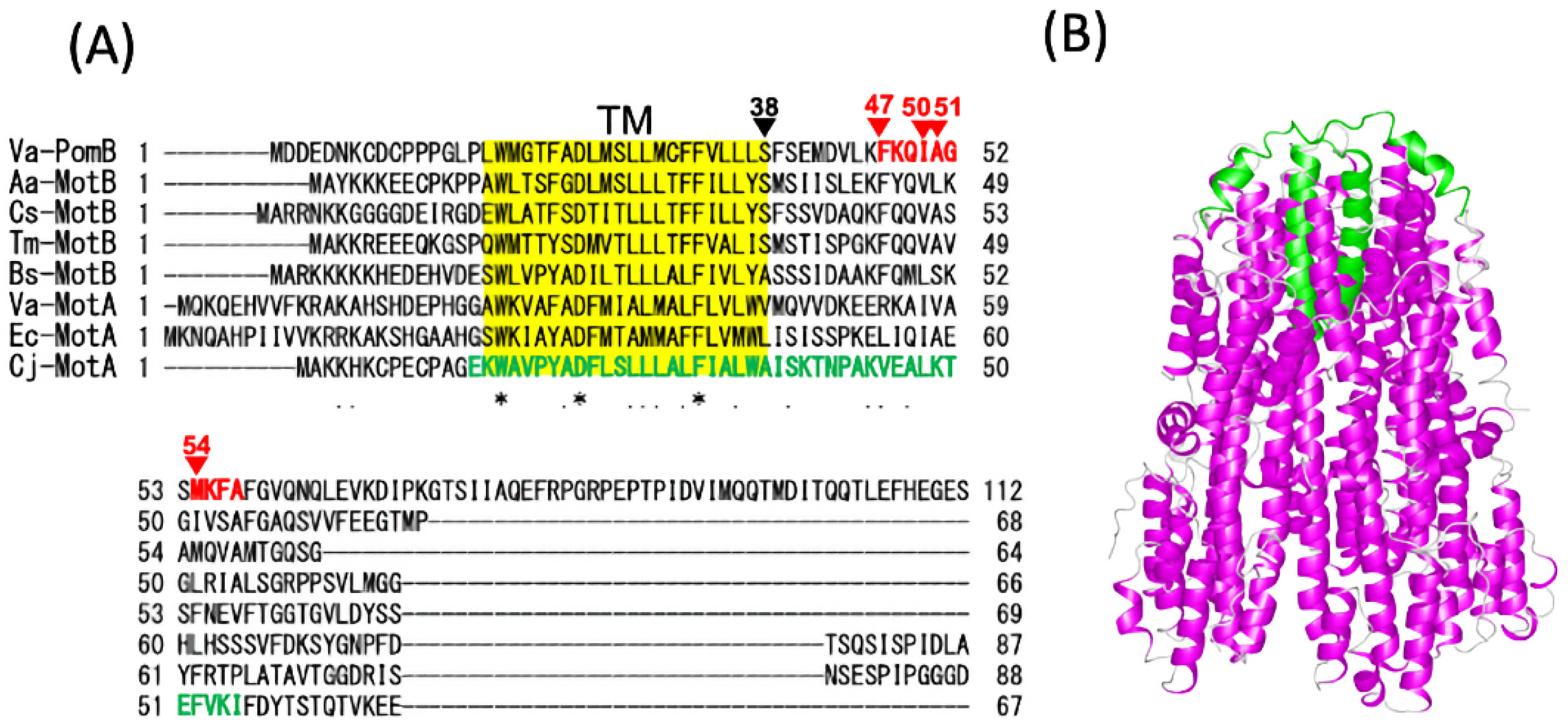
(A) Alignment of the N-terminal sequences of the stator proteins, PomB and MotB. The transmembrane region (TM) is shown in yellow. The residues in the PomB plug region of *Vibrio algynolyticus* (Va) mutated in this study are shown in red. Va: *Vibrio alginolyticus* VIO5, Aa: *Aquifex aeolicus*, Cs: *Clostridium sporogenes*, Tm: *Thermotoga maritima*, Bs: *Bacillus subtilis*, Ec: *Escherichia coli*, Cj: *Campylobacter jejuni*. (B) Atomic structure model of the stator, MotAB, of *C. jejuni* (Cj) (PDB ID: 6YKM). The PomA and PomB parts are shown in magenta and green, respectively.

**Fig. 2.**
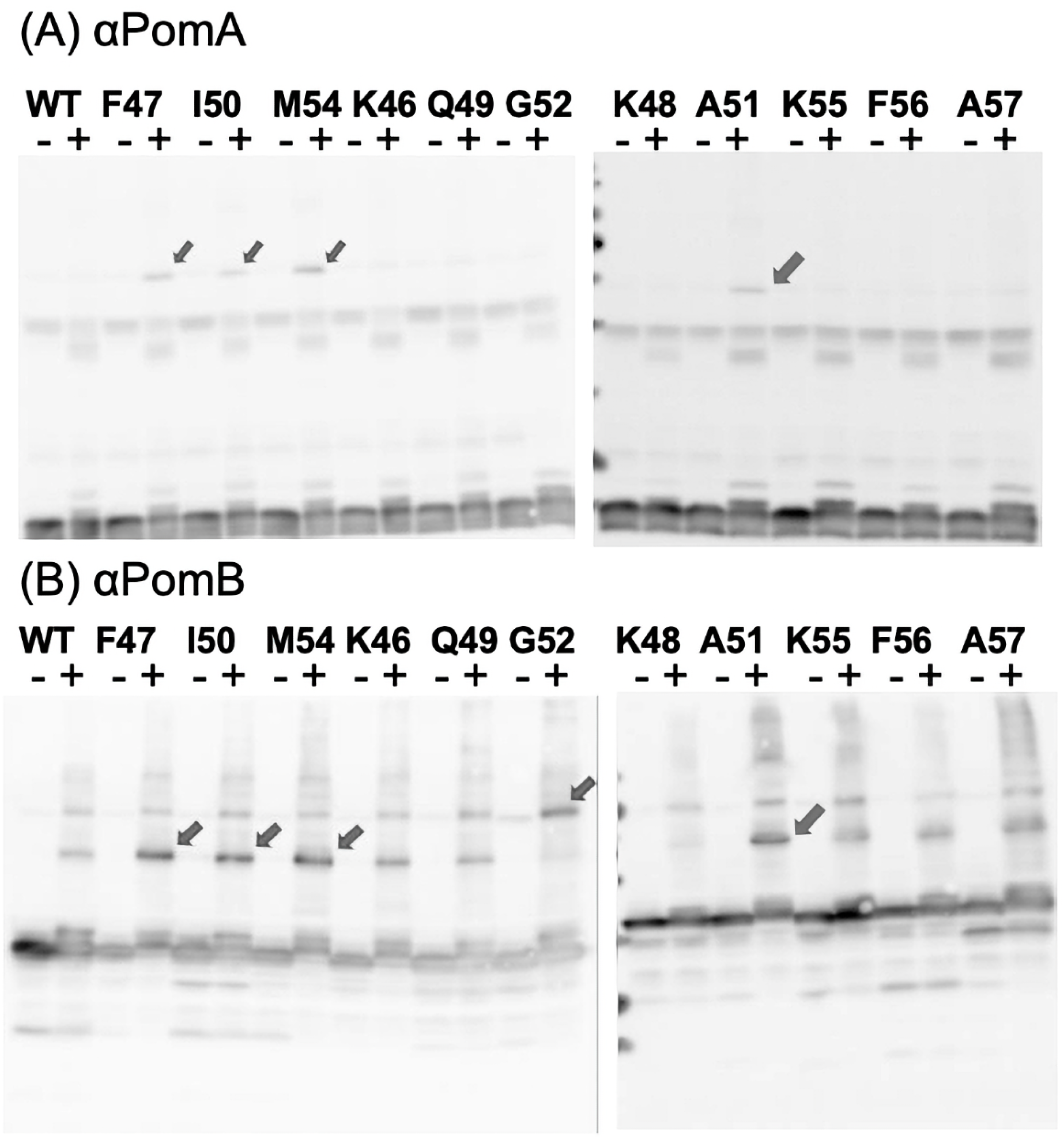
Photo-crosslinking between plasmid-borne *Vibrio* PomA and PomB in *E. coli*. An amber codon was introduced into the position corresponding to each residue of PomB F47 to A57, and the *pomA* and *pomB* genes were expressed in *E. coli* DH5α. At the time, the amber suppressor tRNA was charged with the UV-reactive amino acid, *p*BPA, which is a phenylalanine derivative, to carry out protein synthesis. Proteins extracted from UV-irradiated (+) and non-irradiated (-) cells were separated using SDS-PAGE and detected using western blotting using the anti-PomA antibody (A) and anti-PomB antibody (B).

### Investigation of the interaction between residues of PomA and residues of PomB

To identify the residues of PomA that interact with these residues, we substituted cysteine for the corresponding residues in the structure presumed nearby and examined whether disulfide bonds were formed. Because disulfide bond formation can be analyzed in *Vibrio*, we introduced mutations in the *pomA* and *pomB* genes of the pHFAB plasmid. Of the structure-determined stators, only *C. jejuni* had a structure that included the plug region (27). Although the sequence homology was not high, we could align the residues of PomB-F47, -I50, -A51, and -M54 against the *C. jejuni* residues of MotB-V45, L48, K49, and F52, respectively (Fig. 1). By searching for the corresponding residues of the *Vibrio* PomA residues against the *C. jejuni* residues of MotA, the PomA residues of L166, M169, I175, and G176 were inferred (Fig. S1, S2). The pair mutations with cysteine were introduced into the pHFAB plasmid. As a result, we could detect the bands of increased molecular weight in PomA-M169C and PomB-I50C that reacted with the anti-PomA and anti-PomB antibodies in the absence of reducing agents (Fig. 3AB).

**Fig. 3.**
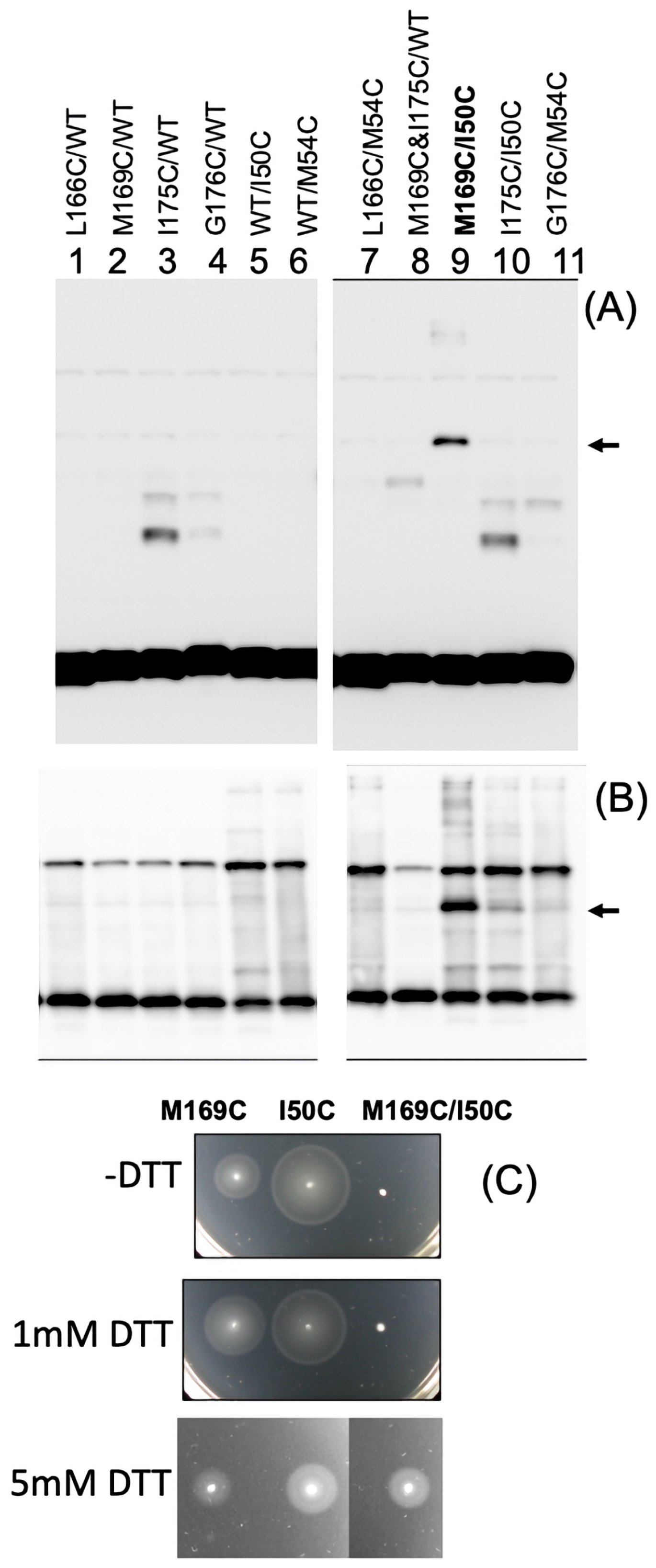
Disulfide-crosslink formation by the introduction of a cysteine residue into the stators, PomA and PomB. Proteins extracted from the *E. coli* DH5α cells harboring the pHFAB-based plasmid with the mutations- 1: *pomA-L166C* and *pomB-wt*, 2: *pomA-M169C* and *pomB-wt*, 3: *pomA-I175C* and *pomB-wt*, 4: *pomA-G176C* and *pomB-wt*, 5: *pomA-wt* and *pomB-I50C*, 6: *pomA-wt* and *pomB-M54C*, 7: *pomA-L166C* and *pomB-M54C*, 8: *pomA-M169C/ I175C* and *pomB*, 9: *pomA-M169C* and *pomB-I50C*, 10: *pomA-I175C* and *pomB-I50C*, and 11: *pomA-G176C* and *pomB-M54C*, were separated using SDS-PAGE in the absence of a reducing agent. PomA and PomB were detected via western blotting using the anti-PomA antibody (A) and anti-PomB antibody (B). Arrows indicate the position of the cross-linked products between PomA and PomB. (C) *Vibrio* NMB191 cells harboring the pHFAB-based plasmid with the mutations-2: *pomA-M169C* and *pomB-wt*, 6: *pomA-wt* and *pomB-M54C*, and 9: *pomA-M169C* and *pomB-I50C*, were inoculated in soft agar plates without DTT and with 1 mM or 5 mM DTT and incubated at 30 °C for 5 hours.

Mutations of PomA-M169C and PomB-I50C were introduced into the *pomAB* gene cloned into the pCold vector, which is capable of large-scale purification (29). On a gel filtration column, the complex was eluted at a position similar to that of the wild-type PomAB complex without the mutations (Fig. S3). By analyzing the non-reducing proteins in that fraction via SDS-PAGE, protein bands that became a ladder with a large molecular weight could be detected by CBB. These proteins are presumed to be cysteine-crosslinked oligomers.

When the *Vibrio* cells producing PomA-M169C or PomB-I50C alone were inoculated into the soft agar medium, the swarm rings formed as wild-type PomA without mutations. This observation indicated that the swarming ability was not inhibited when cysteine was introduced alone. However, in the double mutant, the ability to form swarm rings was completely lost. When the cells were viewed under a microscope, only few motile cells were observed. The double mutant was found to have markedly inhibited stator function. However, the swarm-forming ability was recovered by adding a reducing agent (5 mM DTT) to the medium (Fig. 3C). Therefore, PomA-M169C and PomB-I50C are strongly suggested to be disulfide-bonded by a cysteine residue, while PomA and PomB are bound to each other, resulting in a loss of stator function.

### Ion permeability by mutations in the plug region

In *E. coli*, mutations in the MotB plug regions L55, I58, Y61, and F62 have been shown to completely inhibit growth (20). Based on sequence comparisons, we generated mutants in which the residues F47, I50, and M54 of the PomB plug region corresponding to L55, I58, and F62 were replaced with alanine (A) and glutamic acid (E). The mutants with A showed a phenotype similar to that of the wild-type in motility; however, the mutant with E had a reduced ability to form swarms on soft agar medium (Fig. 4A). Because the decrease in motility due to the mutation in the plug region is known to be caused by growth inhibition (20), the mutants were expressed in *E. coli* and their growth was examined. The overnight culture of *E. coli* was diluted 1/100 and grown. After 3 h, arabinose was subsequently added to express the genes. When the expression of the mutants replaced with A was induced, the growth was similar to that of the wild-type; however, growth inhibition occurred in the mutants replaced with E (Fig. 4B and S4).

**Fig. 4.**
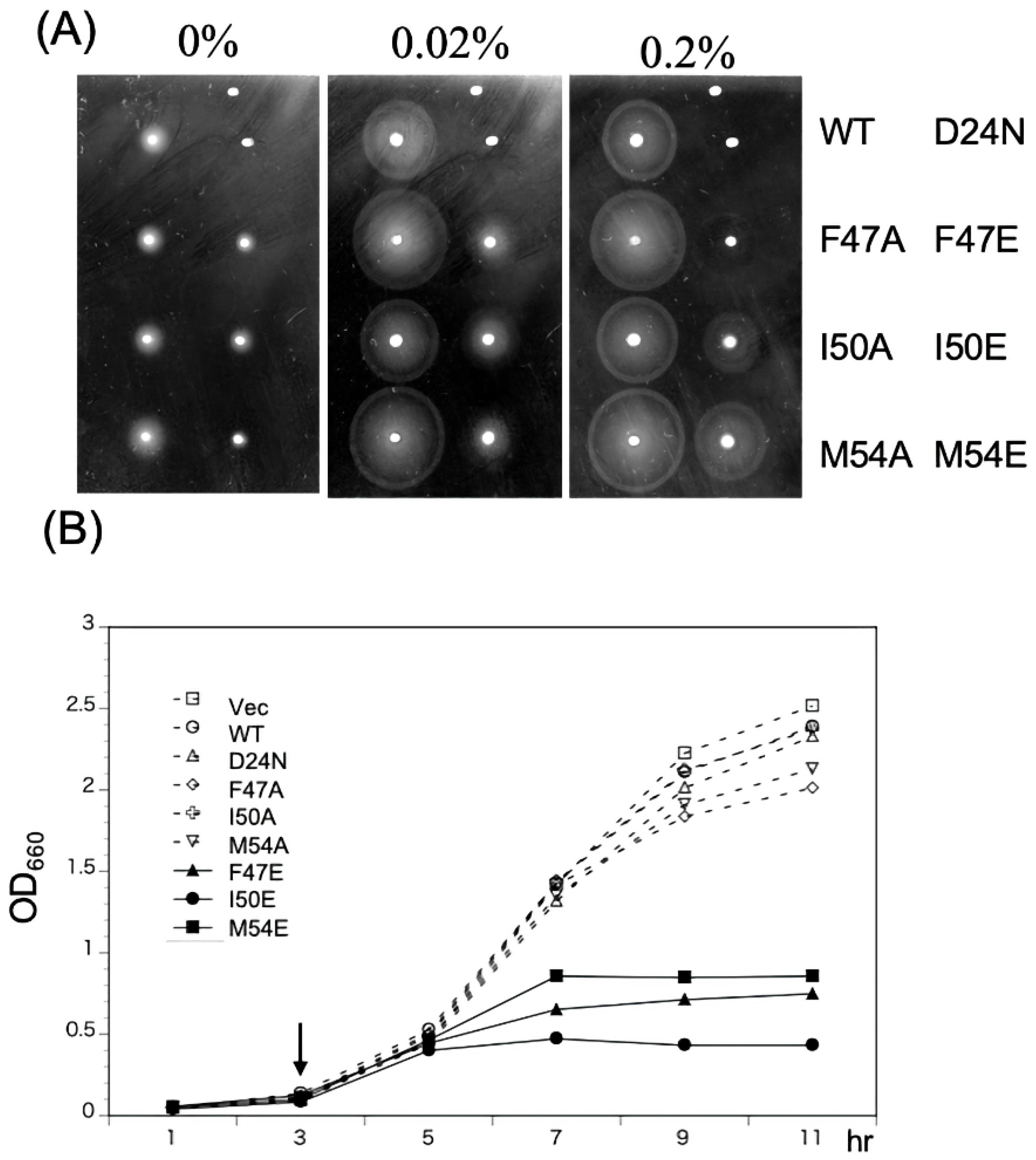
Effect by expression of the PomA/B plug mutant on growth. (A) *Vibrio* NMB191 cells harboring the pHFAB-based plasmid with the mutations, *pomB-wt*, *D24N*, *F47A*, *F47E*, *I50A*, *I50I*, *M54A*, and *M54E*, were inoculated in soft agar plates (VPG 0.25%) with arabinose 0%, 0.02% or 0.2% and incubated at 30 °C for 6 h. (B) Growth curve of cells. Overnight culture of *E. coli* cells harboring the same plasmids as (A) was inoculated into LB 3% NaCl broth at 1/100 dilution; arabinose was added at a final concentration of 0.2% (w/v), 3 hours later, to induce expression (arrow). A_660_ was measured every 2 hours after induction.

### Effects on PomA and PomB cross-link formation by the plug region mutations

We examined the motility of PomA-M169C and PomB-I50C cysteine double mutations (MI), in addition to the PomB-F47E and PomB-M54E mutations. PomB-F47E and PomB-M54E mutations were also introduced into the PomA-D170C and PomB-S38C mutations (DS), which have been previously shown to form a disulfide cross-link (30) (Fig. 5). PomB-F47E and PomB-M54E mutant cells showed reduced swarming ability, which may be attributed to the growth inhibition.

**Fig. 5.**
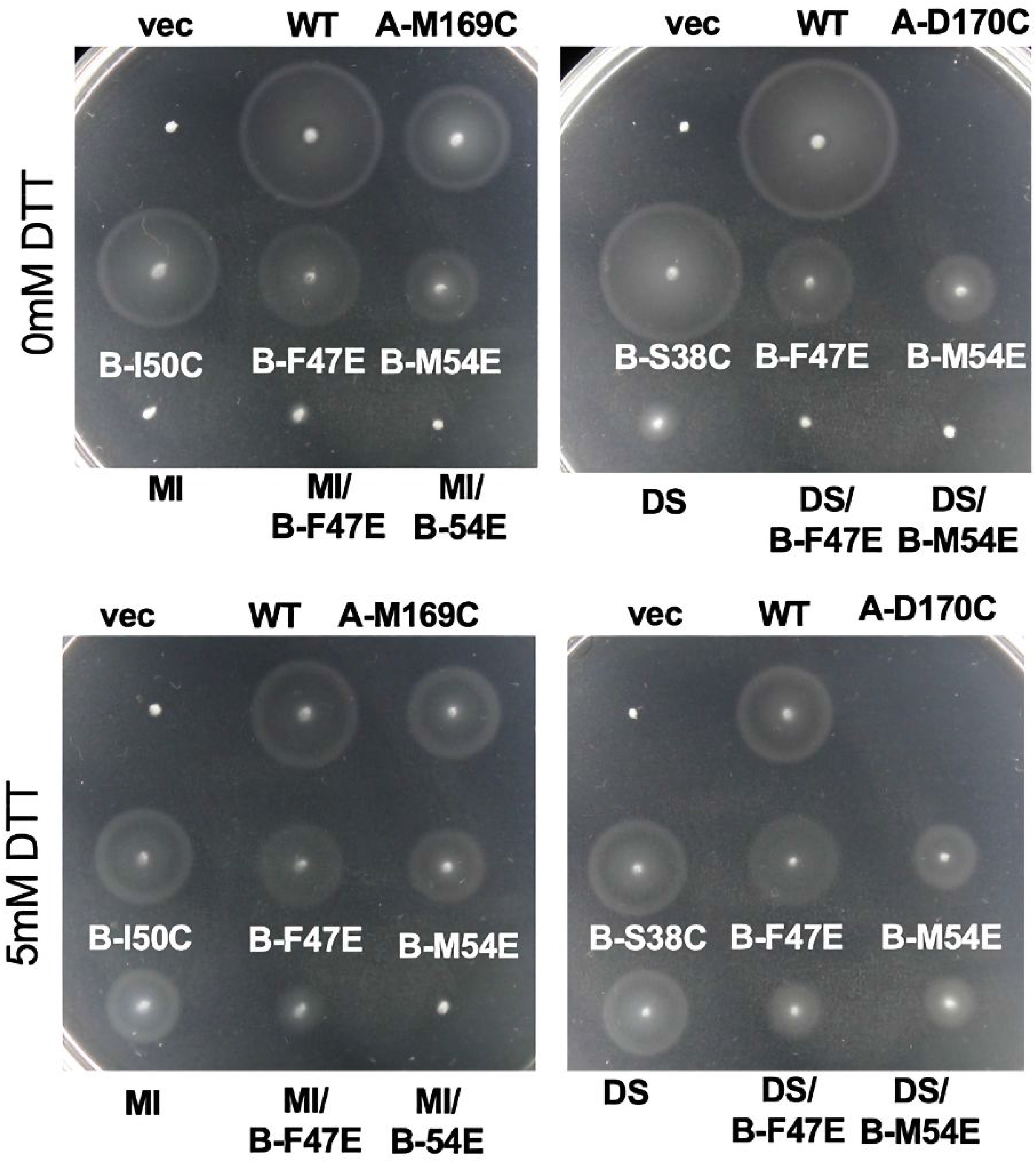
Effect of the plug mutations on the cysteine-substituted mutants. *Vibrio* NMB191 cells harboring pBAD33 (vec), pHFAB (WT), and pHFAB-based plasmid with the mutations, *pomA-M169C* and *pomB-wt* (A-M169C), *pomA-D170C* and *pomB-wt* (A-D170C), *pomA-wt* and *pomB-I50C* (B-I50C), *pomA-wt* and *pomB-F47E* (B-F47E), *pomA-wt* and *pomB-M54E* (B-M54E), *pomA-wt* and *pomB-S38C* (B-S38C), *pomA-M169C* and *pomB-I50C* (MI), MI + *pomB-F47E* (MI/B-47E), MI + *pomB-M54E* (MI/B-54E)*, pomA-D170C* and *pomB-S38C* (DS), DS + *pomB-F47E* (DS/B-47E), and DS + *pomB-M54E* (DS/B-54E) were inoculated in soft agar plates (VPG 0.25%) with arabinose 0.02% and incubated at 30 °C for 5 h with 5 mM DTT or without DTT.

We used SDS-PAGE to examine the crosslinking of MI by introducing PomB-F47E and PomB-M54E mutations. The density of the band at approximately 55 kDa, which is considered the crosslinked band, was reduced by the introduction of the PomB-F47E or PomB-M54E mutation (Fig. 6). The distance between the residues is suggested to be separated by the two mutations, which decreases the reactivity of PomA-M169C and PomB-I50C.

**Fig. 6.**
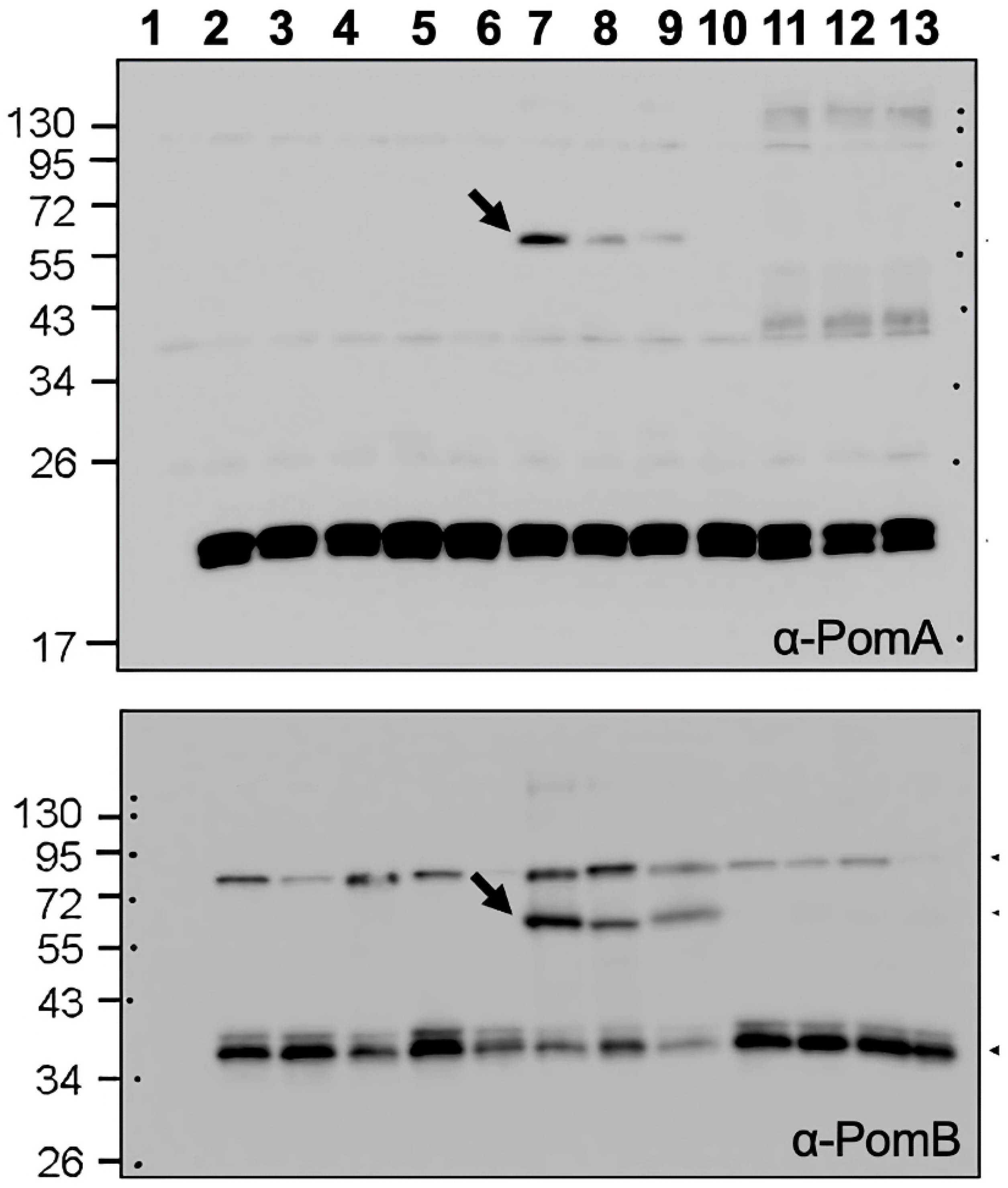
Crosslink formation affected by the plug mutations. *Vibrio* NMB191 cells harboring pBAD33 (1: vec), pHFAB (2: WT/WT), and pHFAB-based plasmid with the mutations-3: *pomA-M169C* and *pomB-wt*, 4: *pomA-wt* and *pomB-I50C*, 5: *pomA-wt* and *pomB-F47E*, 6: *pomA-wt* and *pomB-M54E*, 7: *pomA-M169C* and *pomB-I50C* (MI), 8: MI + *pomB-F47E*, 9: MI + *pomB-M54E*, 10: *pomA-wt* and *pomB-S38C*, 11: *pomA-D170C* and *pomB-S38C* (DS), and 12: DS + *pomB-F47E*, 13: DS + *pomB-M54E*, were grown to the mid-log phase. The proteins of the cells were then separated using SDS-PAGE in the absence of a reducing agent, and detected via western blotting using an anti-PomA antibody (upper) and a PomB antibody (lower). Arrows indicate the position of the cross-linked products between PomA and PomB.

## Discussion

In electrophysiology, the term gating refers to the opening (activation) or closing (deactivation/inactivation) of ion channels. When an ion channel is in the closed state (non-conductive), the ions are impermeable, and the current is not conducted (31). When an ion channel is in the open state, it allows a specific type of ion to pass through it, causing an electric current to flow through the cell membrane. Gating is the process whereby an ion channel transitions between its open and closed states. The mechanism of ion permeation regulation by membrane protein complexes that function in concert with ion permeation is important for living organisms. In general, an ion channel is considered to have four processes: activation, deactivation, inactivation, and reactivation, corresponding to its ionic current state. Its activity is regulated by changes in the voltage applied to the cell membrane, chemicals interacting with the ion channel, temperature, stretching or deformation of the cell membrane, modifications (such as phosphate groups to the ion channel), and interactions with other molecules in the cell (e.g., G-proteins). The ball-and-chain model of voltage-gated channels has been well analyzed in response to ion channels and the mechanisms whereby various cellular changes cause gating (32–34). The mechanism of ion permeation control does not exist only in simple ion channels. In fact, in ATPase, which is a complicated complex, it is known that ε-protein and IF1 regulate ATPase function to regulate ion flow (35–38). Further, owing to the structural transformation of the ε subunit into the F_1_ complex, which is composed of the γ and αβ subunits, the interaction of the ε subunit with the rotation of the γ subunit in the CCW direction is terminated to inhibit ion permeation via ATP degradation. Therefore, under the physiological conditions in most organisms, the CCW rotation of the F_1_F_O_-ATPase is inhibited to avoid energy dissipation due to the wasteful intracellular consumption of ATP.

The mechanism for generating the rotational force in the flagellar motor is thought to be quite different from that of ATPase. However, both are rotational motors that convert ion flow into rotational energy. Until recently, the rotational force of the flagellar motor was thought to be generated by a conformational change in the cytoplasmic region of the stator to push the rotor in a manner similar to the interaction of actin and myosin in the muscle. A structure was reported wherein the transmembrane region of the B subunit in the flagellar stator forms a dimer to become a stalk, and sticks in the center of a pentamer complex formed by the A subunit (27, 28). From this structure, the rotor pentamer was proposed to rotate when the ions flowed between the A and B subunits. Because there are two ion binding sites in the stalk composed of two subunits, 10 steps of 36 ° rotation are predicted to exist in the stator rotation. For the prediction of rotation, a similar situation is seen as the crystal structure had been solved for ATPase and the structure of the asymmetric γ subunit stuck in the center of the αβ trimeric ring where rotation was predicted for the αβ trimeric ring relative to the γ in the F_1_ part (39). In F_1_-ATPase, a famous experiment in which the αβ trimeric ring was fixed to the slide glass revealed the rotation of actin fibers bound to γ (40). The demonstration of the rotation of the flagellar stator between subunits A and B is anticipated.

The flagellar stator was predicted to be a function of ion permeation. Ion permeation was thought to be suppressed when the flagellar motor was not rotated; this is because cell growth was not affected, even when the stator was expressed in large amounts. Although the overexpression of MotA protein was reported to cause some defects in cell growth, it was concluded that the stator protein did not cause ion influx, similar to the ionophores (41, 42). On the other hand, growth inhibition did not occur when MotB was expressed together with MotA. However, growth was inhibited when expressed together with incomplete MotB (43). This growth inhibition is rescued by mutating the ion binding site of the stator complex (44). Based on these studies, it has been hypothesized that the site that inhibits ion influx is located in the periplasmic region of MotB. Growth inhibition was demonstrated to occur at the mutations around residues 52 to 65 of MotB, which are thought to form predicted amphipathic helices, just after the transmembrane region of *E. coli* MotB (20). In the present study, similar amino acid mutations in PomB, a component of the sodium ion channel, were shown to inhibit growth.

The stator complex composed of A and B subunits is assembled in the cytoplasmic membrane; this complex is inactive and does not permeate ions until the stator complex interacts with the rotor. Once the stator comes into contact with the rotor, a structural change in the plug region is induced to release the inhibition, thereby resulting in channel activation. At this time, the periplasmic region of the B subunit is also changed in the structure to interact with peptidoglycan, resulting in anchoring to the cell wall (19, 21). Until now, how ion permeation is blocked and the role of the plug region have been unknown. Although the structures of several stator species have been solved, the structure of the plug region is only visible in *C. jejuni* (27). We assume that the association between the plug region of MotB and the loop region of MotA might be stronger than that of the other species. According to the atomic structure of the stator, a hypothetical model, the stator rotates by the ion flux. Thus, the plug region of PomB must stop or block the rotation between MotA and MotB, rather than cover or block the channel. Structural analysis of *C. jejuni* showed that the loop region (Loop_3-4_), which connects TM3 and TM4 of PomA, is located extracellularly at the inner ring of the PomA pentamer and interacts with the α-helix of the plug region. The structure strongly supports the idea that the plug region stops rotating by trapping it and prevents ion conductance. Our studies have shown that the loss of the plug region does not abolish motor function but increases ion permeability and causes growth inhibition (22, 23). In the present experiments, we found that the plug region interacted with loop_3-4_ to stop ion influx. In addition, our cysteine crosslinking experiments clearly showed that the interaction between the plug and loop regions was weakened by the amino acid mutations that have an inhibitory effect on the growth of the plug region.

The extracellular loop region of *Vibrio* pomA has been analyzed using cysteine scanning mutations (45). Despite the same extracellular loops (i.e., Loop_3-4_ and Loop_1-2_), the profiles of maleimide modification in the Cys mutants were very different; most residues of Loop_3-4_ were modified, while those of Loop_1-2_ were not. At that time, the reason for this difference was unknown. However, when the structure was observed, we understood that Loop_1-2_ was buried in the membrane and it was hard to attach the residues using the maleimide reagent. The cysteine mutants of Loop_1-2_ did not almost affect the motility, except for the D31C mutation. Based on this structure, D31 appears to form an inlet for ions from the membrane surface (46). However, the cysteine mutants of Loop_3-4_ showed a large decrease in swarm size from M169 to K173 of PomA, which was further affected by the cysteine modifier, DTNB. In this study, M169 was shown to interact with the I50 of PomB in the plug region. The PomA-P172C mutant forms a dimer and inhibits its function (47). In the pentamer structure, they are not close enough to form a cross-link. Further, we assume that dimer formation may prevent stator assembly or pentamer formation.

We proceeded to answer the following question: how are the activated and inactivated states regulated? The cytoplasmic loop region (Loop_2-3_) of subunit A in the stator interacts with the C-terminal region of FliG in the C-ring to induce conformational changes (18). The interaction site of FliG and PomA in the cytoplasmic region was recently revealed by photo-crosslinking experiments (48), which were also conducted in this study. We proposed a model in which the sites between FliG and PomA engage and rotate like gears. There are charged residues in the cytoplasmic helices H1, H2, and H3 domains at the cytoplasmic ends of TM3 and TM4, which are important for the interaction with FliG. This domain could thus influence the structure of TM3 and TM4 to change and transmit structural changes to the extracellular region. This structural change would result in the breakage of the interaction between the plug region and the Loop_3-4_ region, thereby resulting in the release of the stopper. Simultaneously, structural changes in the B subunit are expected in the vicinity of the periplasmic region, causing the upright structure suggested by the structural and biochemical analyses (19). A closer look at the structural analysis of *C. jejuni* (27) revealed that the α-helix of the extracellular domain of TM4 of MotA in the stator, which lacks the plug region, is longer than the plug containing the structures. This structural change may lose the flexibility of the coil region connecting the plug region, resulting in the plug region standing up. Either *Vibrio* FliL localizes with the stator complex and the linker region, or the plug region of PomB is necessary for the localization of FliL (49). Although the function of FliL is unclear, it has been shown to be necessary in *Vibrio* cells to produce sufficient torque when the external environment becomes highly viscous (50). It has been speculated that FliL is similar to stromatin, which is present in erythrocytes to regulate ion permeation of transporters and forms a ring structure (51, 52). There is no direct evidence to show that FliL supports the structure of PomB and controls ion permeability. In any case, analysis of the complete stator structure, including the periplasmic region, is required.

## Materials and Methods

### Bacterial strains and plasmids

The bacterial strains and plasmids used in this study are listed in Table S1. *E. coli* was cultured in LB broth (1% (w/v) bactotryptone, 0.5% (w/v) yeast extract, 0.5% (w/v) NaCl], LB 3% NaCl broth [1% (w/v) bactotryptone, 0.5% (w/v) yeast extract, 3% (w/v) NaCl], and TG broth [1% (w/v) bactotryptone, 0.5% [w/v] NaCl, 0.5% [w/v] glycerol). Chloramphenicol was added at a final concentration of 25 μg/mL for *E. coli*. Ampicillin was added at a final concentration of 100 μg/mL for *E. coli*. *V. alginolyticus* was cultured at 30 °C in VC medium (0.5% [w/v] polypeptone, 0.5% [w/v] yeast extract, 0.4% [w/v] K_2_HPO_4_, 3% [w/v] NaCl, 0.2% [w/v] glucose) or VPG medium [1% (w/v) polypeptone, 0.4% (w/v) K_2_HPO_4_, 3% (w/v) NaCl, 0.5% (w/v) glycerol]. If needed, chloramphenicol was added at a final concentration of 2.5 μg mL^−1^ for *V. alginolyticus* culture.

### Photo-crosslinking experiments

*E. coli* DH5α cells harboring two different plasmids, pEVOL-pBpF and pBAD24-based plasmid (pJN142), were cultured in TG broth containing 1 mM *p*-benzoyl-L-phenylalanine (*p*BPA) (Bachem AG, Switzerland), as described previously (48). Cells were collected by centrifugation, suspended in PBS buffer (137 mM NaCl, 2.7 mM KCl, 10 mM Na_2_HPO_4_, 1.76 mM KH_2_PO_4_), collected by centrifugation, and re-suspended in PBS buffer. UV irradiation was carried out with a B-100AP UV lamp (Analytik Jena US, Upland, CA, USA) for 5 min. The cells collected by centrifugation were suspended in sodium dodecyl sulfate (SDS) loading buffer (62.5 mM Tris-HCl [pH 6.8], 2% [w/v] SDS, 10% [w/v] glycerols, 0.01% [w/v] bromophenol blue) containing 5% (v/v) β-mercaptoethanol. Proteins in the samples were separated using SDS-PAGE and the proteins were detected using rabbit anti-PomA or anti-PomB antibodies, as described previously (53).

### Disulfide crosslinking experiment

*E. coli* DH5α cells harboring the pHFAB-based plasmids were cultured in TG broth containing arabinose at a final concentration of 0.02% (w/v) at 30 °C for 5 h from an initial OD_600_ of 0.05. The cells were collected by centrifugation, and the precipitated cells were suspended in SDS-loading buffer without β-mercaptoethanol to detect disulfide crosslinking. The procedure used for SDS-PAGE and immunoblotting was similar to that used in the photo-crosslinking experiment.

### Swimming assay in agar plates

*Vibrio* NMB191 cells harboring pHFAB-based plasmids were plated on VPG plates with antibiotics. A colony of *Vibrio* cells was inoculated onto VPG agar plates [VPG containing 0.25% (w/v) bactoagar and 0.02% (w/v) arabinose] and incubated at 30 °C for the desired time.

#### Purification of the stator complex of PomAB

From the BL21 (DE3) cells carrying the plasmid, pCold4 *pomApomB-his6*, the stator complex was purified according to a previously described procedure (29, Nishikino, 2020 #121).

#### Growth curves

*E. coli* JM109 cells harboring pHFAB plasmids were grown overnight at 37 °C in LB broth containing 25 μg/mL chloramphenicol. The culture was diluted 1:100 in LB 3% NaCl broth and incubated at 30 °C. Arabinose was added at a final concentration of 0.2% (w/v) after 3 h to induce PomA/PomB expression. A_660_ was measured every 2 h after the induction.

## Acknowledgments

We thank Dr. Peter G. Schultz for the kind gift of pEVOL-pBpF for photo-crosslinking,

This work was supported in part by JSPS KAKENHI Grant Numbers 18K19293 (to S.K.), 18K07108 (to H.T.), and 20H03220 (to M.H.).

## Supporting information

Supplementary information associated with this article can be found in the online version on the publisher’s website.

